# Transcriptomic analysis of bone and fibrous tissue morphogenesis during digit tip regeneration in the adult mouse

**DOI:** 10.1101/643361

**Authors:** Feini Qu, Ilan C. Palte, Paul M. Gontarz, Bo Zhang, Farshid Guilak

**Author notes:** corresponding author Correspondence address: Farshid Guilak, Ph.D., Professor of Orthopaedic Surgery, Couch Biomedical Research Bldg., Room 3121, Campus Box 8233, Washington University, Saint Louis, MO 63110.

## Abstract

Humans have limited regenerative potential of musculoskeletal tissues following limb or digit loss. The murine digit has been used to study mammalian regeneration, where stem/progenitor cells (the ‘blastema’) regrow the digit tip after distal, but not proximal, amputation. However, the molecular mechanisms responsible for this response remain to be determined. We hypothesized that regeneration is initiated and maintained by a gene regulatory network that recapitulates aspects of limb development, whereas a non-regenerative response exhibits fibrotic wound healing and minimal bone remodeling. To test these hypotheses, we evaluated the spatiotemporal formation of bone and fibrous tissues after level-dependent amputation of the murine terminal phalanx and quantified the transcriptome of the repair tissue. We show that digit regeneration is a level-dependent and spatiotemporally controlled process, with distal and proximal amputations showing significant differences in gene expression and tissue regrowth over time. Regeneration is characterized by the transient upregulation of genes that direct skeletal system development and limb morphogenesis, including distal Hox genes. By identifying the molecular pathways regulating regeneration, this work will lead to novel therapies that restore complex tissues after injury.

**Summary Statement:** Murine digit tip regeneration after distal amputation is orchestrated through a transient, limb-specific gene network by blastema cells. Proximal amputation activates an alternate transcriptional program that results in scar formation.

## Introduction

Injuries or diseases that lead to the loss of limbs or digits pose important unmet challenges to the medical community. In the United States, an estimated 185,000 people undergo limb amputations annually due to trauma or secondary to diabetes mellitus, dysvascular disease, or malignancy (Owings and Kozak, 1998; Varma et al., 2014). Although prostheses remain the gold standard of care, they remain costly, require extensive rehabilitation, and ultimately fail to restore the full form and function of the limb (Miller et al., 2017). Therefore, strategies that restore the biological composition and structure of the limb have potential to substantially improve patient outcomes. Unfortunately, the regenerative potential of musculoskeletal tissues is restricted, such that healing often culminates in fibrotic scarring. However, there is clinical evidence for digit tip regrowth following injury, especially in young children (Rinkevich et al., 2015). To potentially harness this regenerative capability in humans, the cellular and molecular mechanisms driving regeneration must be better understood.

Towards this end, the murine digit has emerged as an attractive model to study a true regenerative process in adult mammals. After distal resection of the terminal phalanx bone (P3), the distal portion of P3 is first degraded and extruded (Fernando et al., 2011). Afterwards, a mass of hyperproliferative stem/progenitor cells, collectively called the ‘blastema,’ appear at the wound stump to regrow the bone and surrounding fibrous tissues. Lineage tracing studies indicate that blastema cells are germ layer-restricted and derived from local tissues (Lehoczky et al., 2011; Rinkevich et al., 2011), including the bone marrow and periosteum (osteoblast lineage cells) (Lehoczky et al., 2011; Dawson et al., 2018), blood vessels (endothelial cells and pericytes) (Fernando et al., 2011; Rinkevich et al., 2011), dermis (fibroblasts) (Marrero et al., 2017; Wu et al., 2013), nail matrix (nail stem cells) (Lehoczky and Tabin, 2015), and nerve (dedifferentiated Schwann cells) (Johnston et al., 2016). Remarkably, removing the distal portion of P3 results in the restoration of lost tissues, whereas increasingly proximal amputations lead to scar formation (Neufeld and Zhao, 1995; Chamberlain et al., 2017; Takeo et al., 2013; Dawson et al., 2017). A similar phenomenon occurs after damage to the human fingertip (Rinkevich et al., 2015), making these studies directly relevant to clinical finger injuries.

Despite recent advances in our understanding of the role of the digit blastema in regeneration, it remains to be determined how these diverse cell types and signals work in concert to restore the amputated tissues. The mechanisms responsible for level-dependent healing (or lack of healing) also remain to be determined. A critical anatomical boundary that permits regeneration versus fibrotic scarring may exist, but whether this threshold is determined by the availability of stem/progenitor cell populations, the activation of pro-regenerative genes, and/or the persistence of pro-fibrotic pathways is currently unknown. To probe these questions, previous studies have relied on methods such as *in situ* hybridization, polymerase chain reaction, and microarrays for transcriptomic analysis of the regenerating tissue (Chadwick et al., 2007; Cheng et al., 2013; Johnston et al., 2016), but these methods are limited to quantifying known transcripts. In contrast, next-generation RNA sequencing (RNA-seq) technology enables the discovery of novel and rare transcripts, including non-coding RNAs, with higher specificity and sensitivity and at a wider dynamic range.

Therefore, to investigate the signaling pathways of the digit blastema, we used RNA-seq to determine the transcriptomic response of these cells throughout the time course of regeneration. We hypothesized that successful regeneration depends on the coordinated remodeling and regrowth of bone and adjacent tissues, which is initiated and maintained by a gene regulatory network that recapitulates aspects of limb development. To test this hypothesis, we first quantified the spatiotemporal changes to bone and fibrous tissues after distal amputation (Regenerative; Regen) or proximal amputation (Non-Regenerative; Non-Regen) of P3 using microcomputed tomography (microCT) and multiple histologic and imaging modalities. Next, we quantified the differentially expressed genes of the Regen and Non-Regen repair tissue at multiple time points using RNA-seq. From this data, we identified key transcription factors that may be critical for regeneration and confirmed their expression in the digit tip using RNA fluorescence *in situ* hybridization. Our findings suggest that regeneration may depend on the activation of a limb-specific developmental program, whereas regenerative failure may result from reduced stem/progenitor cell populations and developmental signals, and/or accelerated fibrosis.

## Results

### Temporal dynamics of bone and soft tissue regrowth

Pre- and post-operative microCT imaging revealed that 24.4±2.8% and 64.2±2.1% of the terminal phalanx bone (P3) length was amputated for the Regenerative (Regen) and Non-Regenerative (Non-Regen) groups, respectively (**Fig. 1**). Due to the tapered shape of P3, 7.7±1.0% and 37.5±3.9% of the bone volume was removed for the Regen and Non-Regen groups, respectively. Tracking the morphologic time course of P3 after amputation, we found that resecting the distal portion consistently led to a regenerative response with blastema formation and osteogenesis, whereas proximal amputations resulted in fibrotic scar formation. Both P3 length and volume were reduced for the Regen group after distal amputation at 10 days post-amputation (DPA) and at subsequent time points up to 21 DPA when compared to post-surgical values at 0 DPA, indicating histolysis and tissue remodeling (*P*<0.05, **Fig. 1B, Fig. 1C**). At 10 DPA, Hematoxylin and Eosin (H&E) staining showed incomplete epithelization at the distal tip (**Fig. 2A**). Between 10 and 12 DPA, a distal bone fragment was extruded as the wound epidermis closed over the P3 surface (**Fig. 1A**). Maximum bone volume loss occurred at 12 DPA for Regen digits (48.3±5.5%). This initial stage of partial bone degradation opened the marrow cavity and was followed by a local surge in cellularity and extracellular matrix (ECM) deposition. By 14 DPA, a hypercellular mass (the blastema) extended from inside the marrow cavity to the digit tip (**Fig. 2A**). There was a marked periosteal reaction in Regen digits, most notably along the dorsal surface. By 21 DPA, Picrosirius Red (PSR) staining revealed collagen deposition (**Fig. 2B**) and mineralization (**Fig. 1A**) contiguous with the edges of P3, indicating new bone formation distal to the degraded bone. By 28 DPA, both P3 length and volume had increased to reach post-amputation levels at 0 DPA (*P*>0.05, **Fig. 1B, Fig. 1C**). By the 56 DPA end time point, P3 length had recovered to pre-amputated levels in Regen digits, with a 23% volume overshoot (*P*<0.05, **Fig. 1B, Fig. 1C**), and digit morphology resembled that of unamputated control digits (**Fig. S1**).

**Fig. 1.**
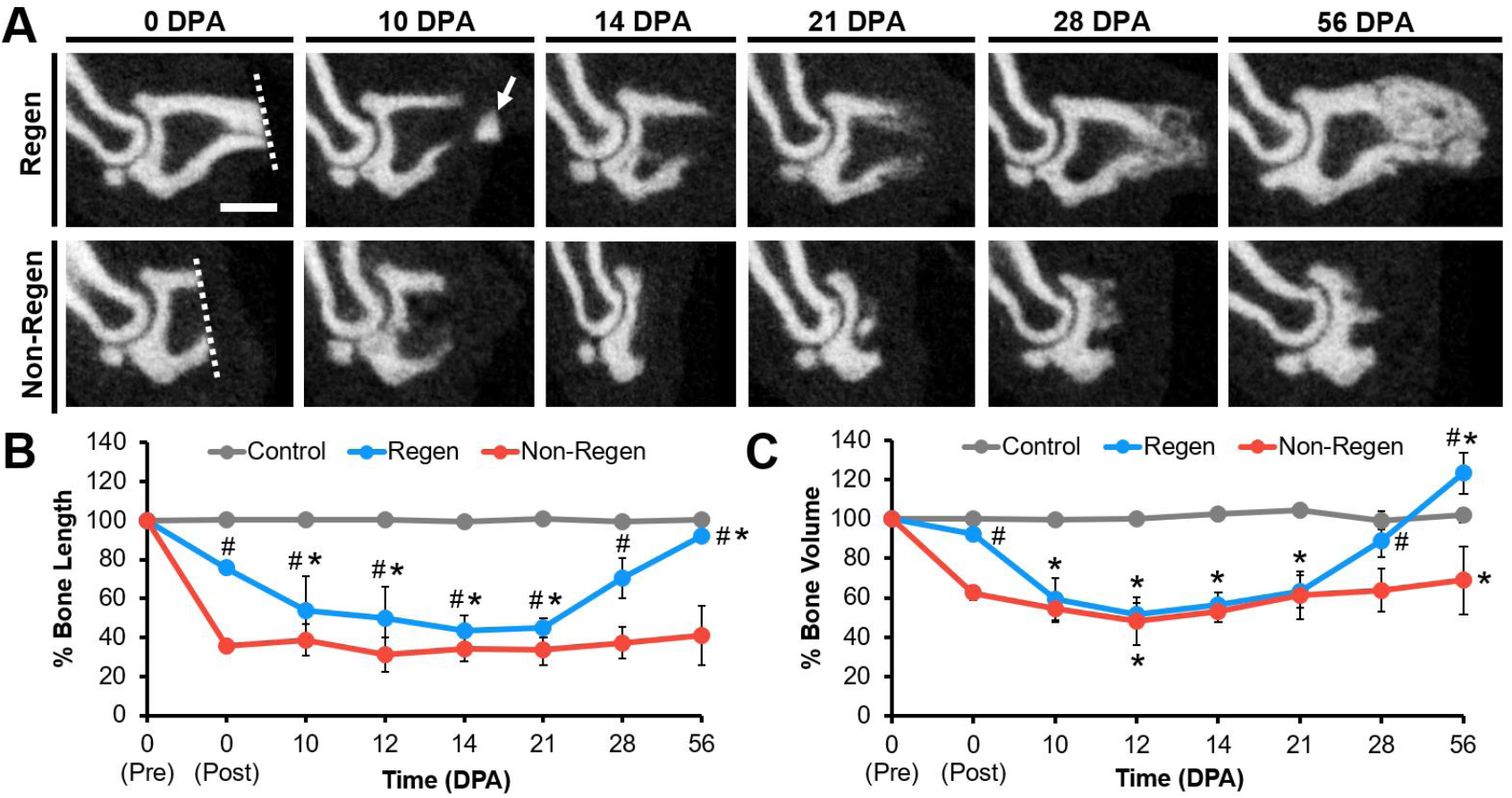
Bone remodeling and regrowth follow distal, but not proximal, amputation. (**A**) Mid-sagittal microCT images of the terminal phalanx bone (P3) at various days post-amputation (DPA) for Regen and Non-Regen groups. Dotted line shows amputation plane. Arrow points to distal fragment. Scale: 0.5 mm. (**B**) P3 length and (**C**) volume changes over time (% of pre-amputation, n=4 mice, 6 digits/group/time point, mean±s.d.). 2-way ANOVA with Tukey’s post-hoc test. \**P*<0.05 vs. 0 DPA post-amputation. ^#^*P*<0.05 vs. Non-Regen.

**Fig. 2.**
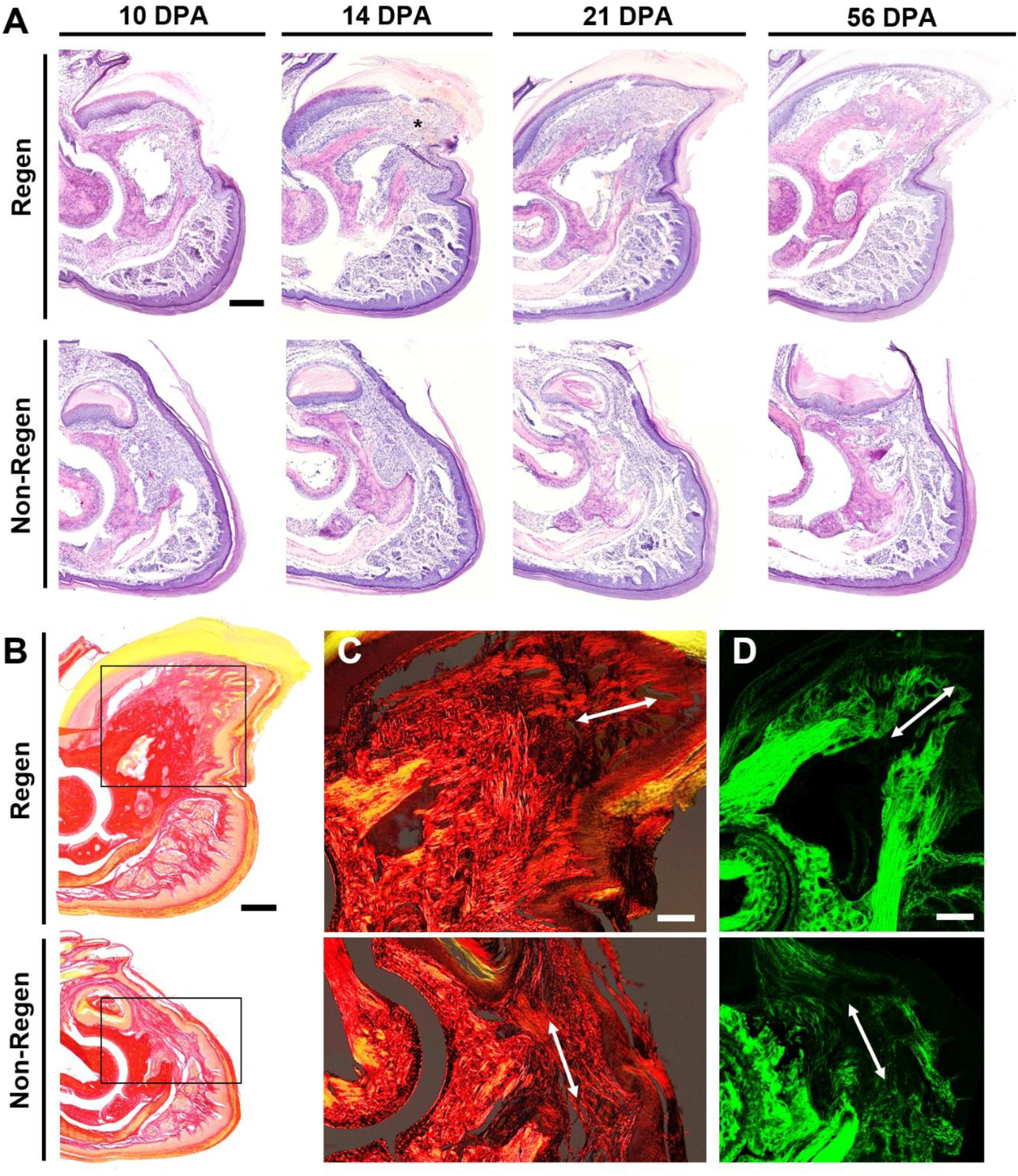
Histological time course of regeneration. (**A**) H&E staining at various DPA for Regen and Non-Regen groups. Asterisk indicates blastema at 14 DPA. Scale: 0.2 mm. (**B**) Picrosirius Red (PSR) staining imaged with bright field microscopy at 21 DPA. Scale: 0.2 mm. Inset shows area imaged in (C). (**C**) PSR staining visualized with polarized light microscopy and (**D**) second harmonic generation (SHG) imaging show collagen fibers (arrow) extending proximodistally in the Regen but not the Non-Regen state at 21 DPA. Scale: 0.1 mm.

Conversely, bone remodeling and regrowth were not observed after proximal amputation in the Non-Regen digits. While P3 volume was slightly reduced at 12 DPA compared to immediately post-surgery at 0 DPA (*P*<0.05, **Fig. 1B, Fig. 1C**), bone length remained constant through 56 DPA (*P*>0.05). Cell accumulation at the cortical bone stump and complete epithelization was apparent by 10 DPA, but minimal periosteal reaction, cellular outgrowth, or bone regeneration was observed at subsequent time points (**Fig. 2A**). Instead, collagen deposition at the endosteal surface was apparent by 14 DPA, and by 21 DPA, the bone stump, as well as the overlying nail matrix and nail plate, was surrounded by a thin fibrous capsule that became more pronounced over time (**Fig. 2B**). At 56 DPA, this conformation remained mostly unchanged, despite continued growth of the nail plate. P3 length and volume did not significantly change for control digits over the course of the study. Furthermore, the single and double-digit amputation models behaved similarly, suggesting that the healing response acts locally. This initial data established the feasibility and timeline of the murine digit amputation model for probing regenerative success versus failure.

### Collagen microstructure and alignment reflect tissue outgrowth

To examine the ECM structure of amputated digits, we utilized polarized light microscopy to visualize collagen distribution and alignment in PSR-stained tissue sections (**Fig. 2C, Fig. S2A**). Over time, collagen at the distal tip became increasingly aligned in the proximodistal direction for Regen digits and in the dorsoventral direction for Non-Regen digits. These findings were confirmed with second harmonic generation (SHG) imaging (**Fig. 2D, Fig. S2B**), which revealed fine collagen fibrils emanating from the bone stump of Regen digits by 14 DPA, converging to form the tip of P3 by 21 DPA. In contrast, a dense network of thick collagen bundles traversed parallel to the amputation plane in Non-Regen digits by 14 DPA. Collectively, these results point to the markedly disparate healing trajectories in response to level-dependent amputation of the same bone.

### Activation of limb-specific developmental processes after distal amputation

Given these morphologic observations, we next examined the molecular signals that delineate regenerative success (Regen) versus failure (Non-Regen) using RNA-seq at time points that reflect blastema formation (12 DPA) and differentiation (14 DPA), and the beginning of bone regrowth (21 DPA). Principal component analysis (PCA) revealed sample clustering by group and time point (**Fig. 3A**). While the Regen and Non-Regen groups remained largely separate, Non-Regen digits at 14 DPA overlapped with Regen digits at 12 DPA. Pairwise comparisons between time points within each group revealed 1513 and 1400 differentially expressed genes (DEGs) in Regen and Non-Regen digits, respectively, the majority of which were protein coding transcripts (**Fig. S3, Table S1**). DEGs were subsequently grouped into 4 clusters based on the time course of expression using k-means clustering (**Fig. 3B**). Although DEGs within each cluster exhibited similar fold changes for Regen and Non-Regen groups, only a quarter of the 1513 Regen DEGs had the same expression pattern in the Non-Regen group, suggesting distinct transcriptomic differences between Regen and Non-Regen digits (**Fig. 3C**). Furthermore, the majority of Regen DEGs exhibited transient upregulation at 14 DPA (49.8%), whereas the majority of Non-Regen DEGs showed downregulation with time (38.9%).

**Fig. 3.**
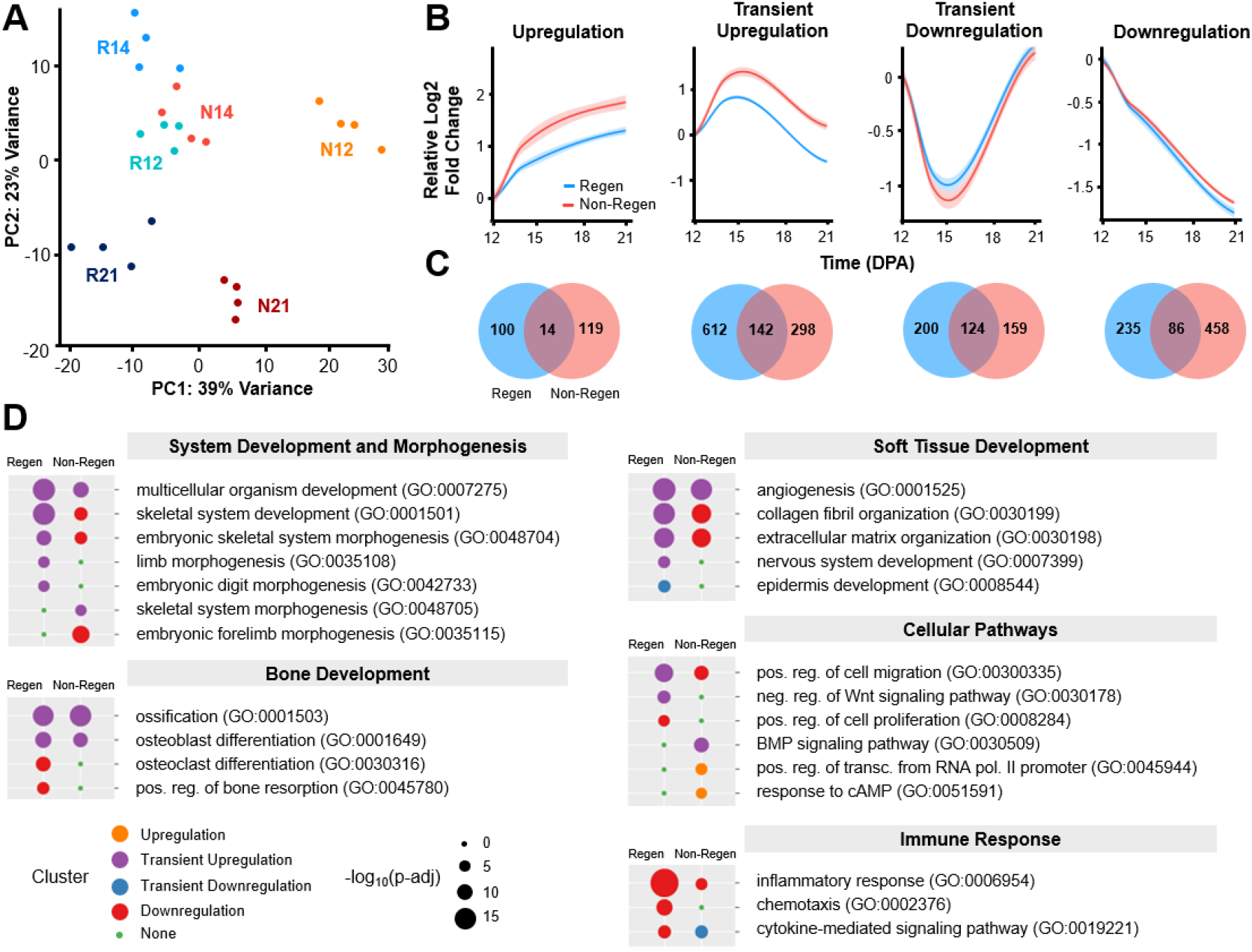
Temporal transcriptomic dynamics of regeneration success versus failure. (**A**) Principle component analysis (PCA) illustrates the relative similarity between Regen (R) and Non-Regen (N) groups at 12, 14, and 21 DPA (n=4 mice, 4 digits/group/time point). (**B**) K-means clustering identified four gene clusters with similar expression patterns. Solid lines represent the mean relative log2 fold change in expression level (RPKM) compared to 12 DPA. Lighter shades indicate 95% confidence interval. (**C**) Venn diagrams show number of genes present in Regen, Non-Regen, or both groups for each cluster. (**D**) Relevant GO terms (biological processes) for Regen and Non-Regen groups, where the circle color and size represent cluster type and Benjamini corrected p-value (p-adj), respectively.

Gene ontology (GO) enrichment analysis revealed that biological processes related to digit regeneration, including skeletal system development, embryonic skeletal system morphogenesis, limb morphogenesis, and embryonic digit morphogenesis, were transiently upregulated in Regen digits but not in Non-Regen digits (Benjamini corrected *P*<0.05, **Fig. 3D, Table S2**). Genes in the Regen transient upregulation cluster also identified with pro-regenerative terms such as multicellular organism development, collagen fibril and ECM organization, osteoblast differentiation and ossification, angiogenesis, and nervous system development. Transiently upregulated cellular pathways include positive regulation of cell migration and negative regulation of the Wnt signaling pathway. Epidermis development fell in the transient downregulation cluster, most likely due to the comparative upregulation of other processes at 14 DPA. The Regen downregulation cluster contained genes related to bone remodeling (osteoclast differentiation and bone resorption), cell division (positive regulation of cell proliferation), and immune response (inflammatory response, chemotaxis, and cytokine-mediated signaling pathway), indicating that these processes had peaked by 12 DPA.

In contrast, while Non-Regen digits shared a subset of transiently upregulated developmental GO terms (multicellular organism development, ossification, osteoblast development, and angiogenesis) (Benjamini corrected *P*<0.05), limb morphogenic processes either fell in the downregulation cluster or were absent (**Fig. 3D, Table S2**). Biological processes related to bone degradation and soft tissue development (nervous system and epidermis) were also absent in Non-Regen digits. Indeed, out of the 754 transiently upregulated Regen DEGs (**Fig. 3C, Fig. 4A**), including 60 transcription factors (TFs) (**Table S1**), only 142 of those same genes (13 TFs) were also transiently upregulated in Non-Regen digits at 14 DPA. Interestingly, the bone morphogenetic protein (BMP) signaling pathway was transiently upregulated, suggesting that there may be a pro-osteogenic response at 14 DPA at the transcriptomic level. These data suggest that although Regen and Non-Regen digits may share some similar signaling pathways, the temporal activation of a limb-specific developmental program may be necessary for a fully regenerative outcome.

**Fig. 4.**
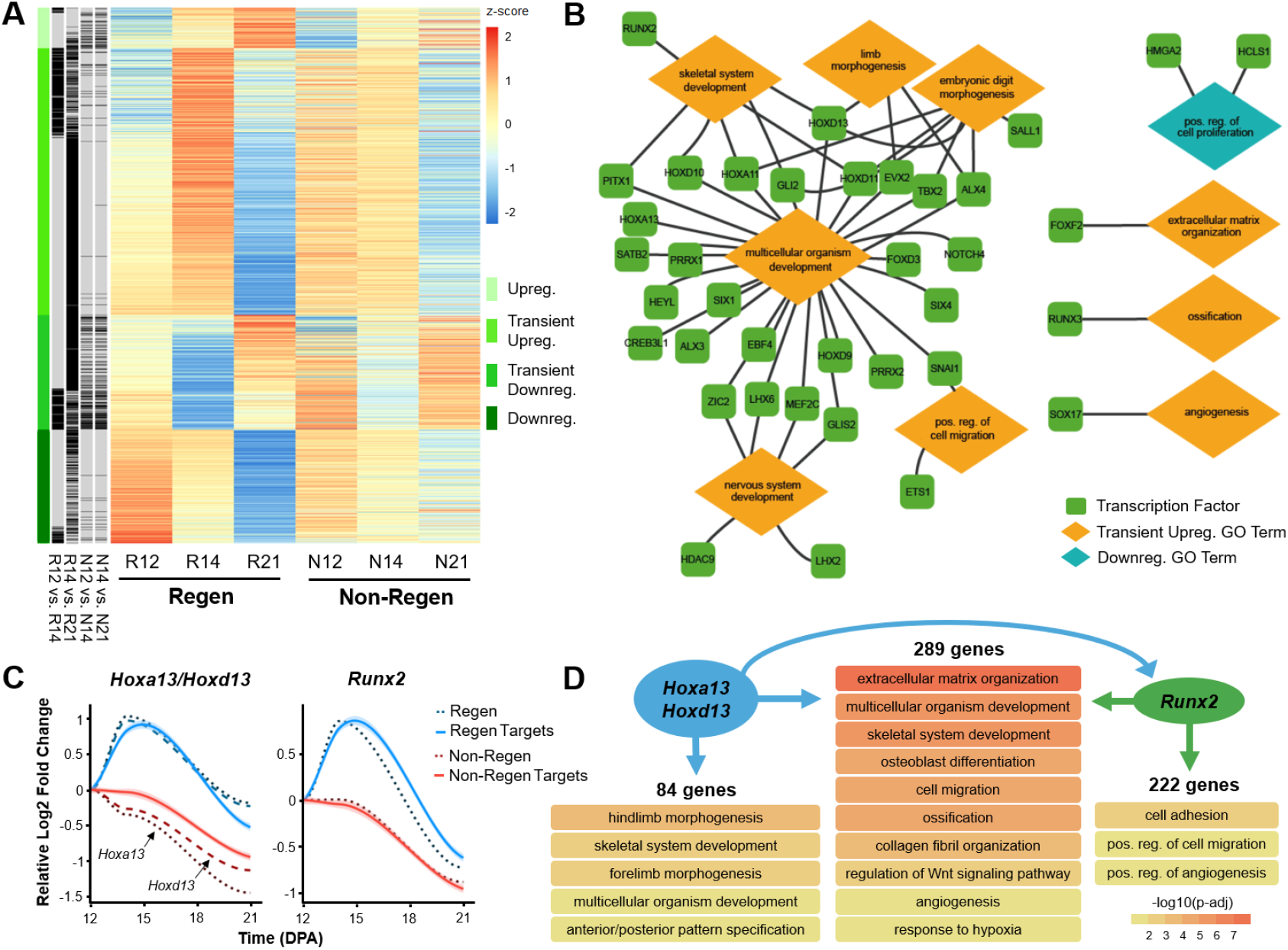
Developmental transcriptional program is transiently activated in regenerating digits. (**A**) Heat map shows the scaled z-score of the average RPKM of 1513 differentially expressed genes (DEGs) from Regen groups at 12, 14, and 21 DPA, alongside the corresponding z-score of Non-Regen groups. Red and blue cells indicate relative gene upregulation and downregulation, respectively. Green cells indicate cluster type, and black cells indicate genes that are significantly different between time points for Regen and Non-Regen groups (Benjamini corrected *P*<0.05). (**B**) Gene regulatory network illustrates various Regen transcription factors (TFs) and their relationship to Regen GO terms. Lines indicate association of TF with GO term. (**C**) Gene expression profiles of the TFs *Hoxa13*/*Hoxd13* (Hox13) and *Runx2* (dashed lines) and their downstream targets (solid lines), as determined by ChIP-seq analysis (n=4 mice, 4 digits/group/time point, mean). Lines represent the relative log2 fold change in expression (RPKM) compared to 12 DPA. Lighter shades indicate 95% confidence interval. (**D**) GO terms (biological processes) associated with gene targets of either Hox13, *Runx2*, or both (Benjamini corrected *P*<0.05). Arrows specify direction of gene regulation, including Hox13 regulation of *Runx2* expression.

### Gene regulatory network reveals TFs driving regeneration

To further investigate how specific genes may direct regeneration, we built a gene regulatory network (GRN) using 37 Regen TFs that were related to pro-regenerative GO terms (2 downregulated TFs and 35 transiently upregulated TFs, 10 of which were also transiently upregulated in Non-Regen digits) (**Fig. 4B, Fig. S4**). Most TFs were associated with multicellular organism development and may also be linked to skeletal system development, limb morphogenesis, embryonic digit morphogenesis, nervous system development, and/or positive regulation of cell migration. TFs of note include Hox genes, which direct distal limb (*Hoxa11*/*Hoxd11*) and digit development (*Hoxa13*/*Hoxd13*) (Fromental-Ramain et al., 1996; Villavicencio-Lorini et al., 2010; Zakany and Duboule, 1999); *Prrx1*, which is required for early skeletogenesis and modulates limb segment length (Cretekos et al., 2008; ten Berge et al., 1998); *Alx4* and *Tbx2*, which affect limb patterning (Kuijper et al., 2005; Sheeba and Logan, 2017); and *Runx2*, which is a master regulator of osteoblast differentiation (Komori, 2010). Other transiently upregulated TFs in Regen digits include those involved in ECM organization (*Foxf2*) (Nik et al., 2016), ossification (*Runx3*) (Bauer et al., 2015), and angiogenesis (*Sox17*) (Corada et al., 2013). One downregulated TF linked to positive regulation of cell proliferation, *Hmga2*, is associated with stem cell self-renewal (Hammond and Sharpless, 2008) and tumorigenesis in various tissues (Pallante et al., 2015).

Next, to determine the downstream implications of transient TF upregulation, we analyzed the DNA binding targets of *Hoxa13/Hoxd13* (hereafter referred to as Hox13) (Sheth et al., 2016) and *Runx2* (Meyer et al., 2014) using publicly available chromatin immunoprecipitation followed by high-throughput sequencing (ChIP-seq) data. We identified 373 target genes of Hox13 and 511 target genes of *Runx2* that were also transiently upregulated in Regen digits, 289 of which were shared targets of Hox13 and *Runx2* (**Fig. 4C**). GO enrichment analysis revealed that unique targets of Hox13 were specifically related to limb morphogenesis, whereas shared targets were associated with processes involved in osteogenesis (e.g., ECM organization, osteoblast differentiation, ossification, and angiogenesis) (**Fig. 4D, Table S3**). Furthermore, the ChIP-seq data revealed that Hox13 could regulate *Runx2* by binding 5.2 kb upstream of a major intragenic transcription start site (**Fig. S5**). While the data did not show direct regulation of Hox13 by *Runx2, Runx2* may potentially affect the transcription of *Hoxd13*, but not *Hoxa13*, via an upstream enhancer. The same target genes in Non-Regen digits reflected the downregulation of Hox13 and *Runx2* in those samples (**Fig. 4C**), which is consistent with our analysis.

### Spatiotemporal expression of regulatory genes

To validate RNA-seq results and to localize gene expression, we performed RNA fluorescence *in situ* hybridization (FISH) for two transiently upregulated TFs, *Hoxa13* and *Runx2*, as well as for *H19*, a long non-coding RNA (lncRNA) that showed significantly higher expression in Non-Regen digits compared to Regen digits at 12 and 14 DPA (**Fig. 5A**). RNA FISH showed increased expression of both *Hoxa13* and *Runx2* in Regen digits compared to Non-Regen digits at 14 DPA (**Fig. 5B**), reflecting the transcriptomic data. In Regen digits at 14 DPA, the blastema (cell mass distal to P3), bone interface, and distal portions of the marrow cavity and endosteum showed areas of high *Hoxa13* and *Runx2* expression, whereas the distal periosteum was largely restricted to *Runx2* expression (**Fig. 5C**). *Hoxa13* and *Runx2* were expressed at comparatively lower levels in Regen digits at 12 DPA (**Fig. S6**), suggesting that their expression significantly increased between the stages of blastema formation and differentiation. The lncRNA *H19* was found in Non-Regen digits at all time points, predominantly located at the bone interface and within the marrow cavity (**Fig. 5, Fig. S6**). *H19* was minimally expressed in Regen digits at 12 and 14 DPA but was present in the region between the distal regenerating bone and epidermis at 21 DPA. *H19* was also concentrated at the ventral epidermal-dermal junction in all samples, including control digits (**Fig. S1**). Taken together, these findings highlight the dynamic, spatiotemporal gene expression patterns that occur after digit amputation, resulting in signaling pathways that may either promote or inhibit the regenerative process.

**Fig. 5.**
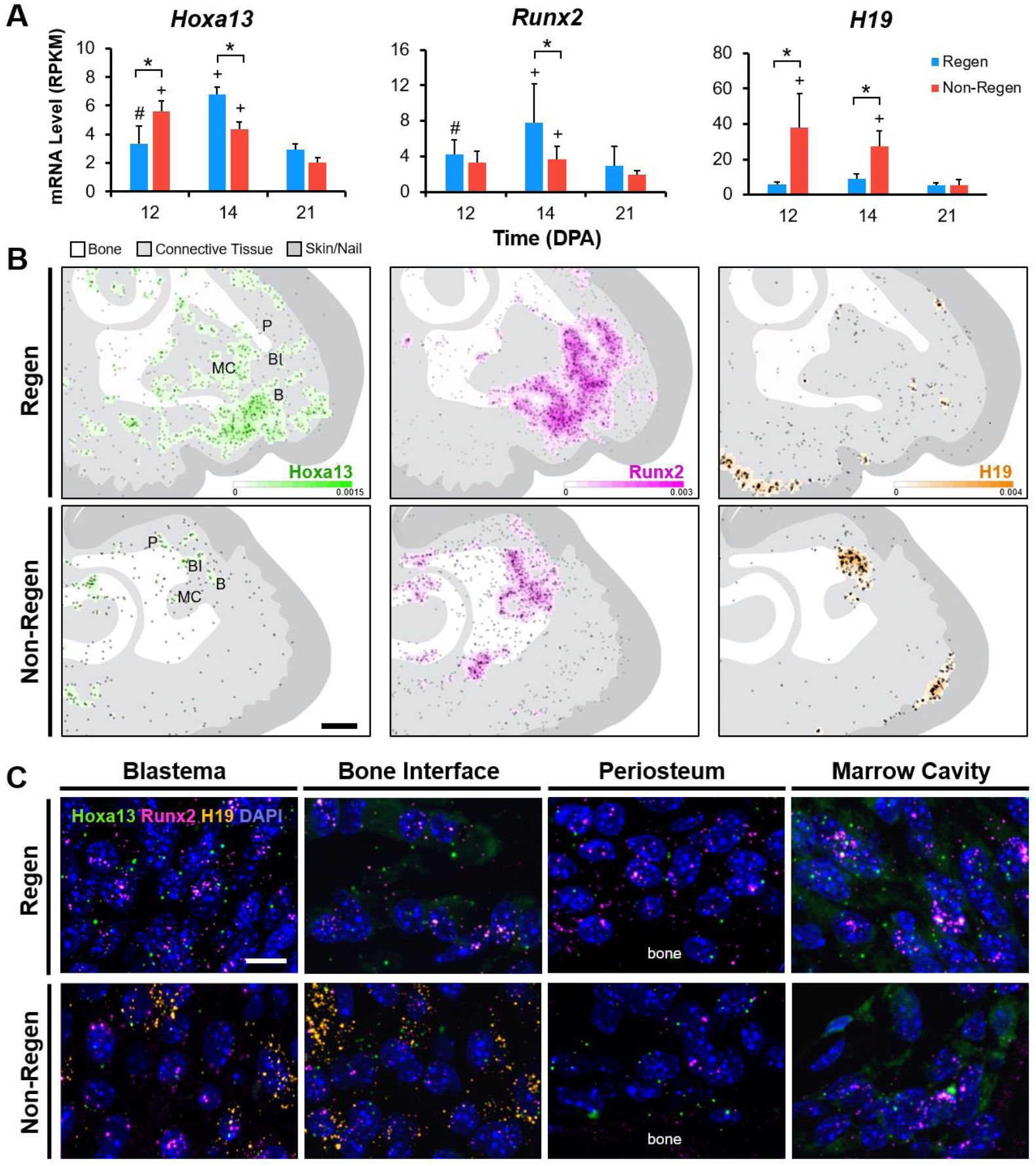
Gene expression is localized to regenerating tissues. (**A**) Bulk RNA-seq mRNA levels (RPKM) of *Hoxa13, Runx2*, and *H19* (n=4 mice, 4 digits/group/time point, mean±s.d.). 2-way ANOVA with Tukey’s post-hoc test. \**P*<0.05 vs. Non-Regen. ^#^*P*<0.05 vs. 14 DPA. ^+^*P*<0.05 vs. 21 DPA. (**B**) Density heat maps show estimated RNA FISH probes per unit area for *Hoxa13* (green), *Runx2* (magenta), and *H19* (orange) in Regen and Non-Regen digits at 14 DPA. Circles represent individual probe signals. Regions of interest shown in (C) include the blastema (B), bone interface (BI), periosteum (P), and marrow cavity (MC). Scale: 0.2 mm. (**C**) Confocal images at 63X magnification show RNA FISH probes for *Hoxa13, Runx2*, and *H19*, and cell nuclei (DAPI, blue) in various Regen and Non-Regen tissues at 14 DPA. Scale: 10 µm.

## Discussion

This work established the transcriptomic landscape of bone and fibrous tissue remodeling and regrowth in the murine P3 amputation model for probing musculoskeletal regeneration. We found that digit tip regeneration is a level-dependent and spatiotemporally controlled process, with distal amputation (Regen) and proximal amputation (Non-Regen) groups showing significant differences in bone remodeling, ECM organization, and gene expression over time (**Fig. 6**). Regen digits are characterized by a limb-specific transcriptional program that result in aligned collagen deposition and ossification extending distally from the amputated cortical bone stump. Interestingly, despite the presence of numerous genes related to osteogenesis, Non-Regen digits did not exhibit significant bone regrowth, but instead produced a fibrous capsule that may hinder tissue outgrowth. These novel findings support our hypothesis that successful regeneration depends on the temporal activation of a gene regulatory network that recapitulates aspects of limb development.

**Fig. 6.**
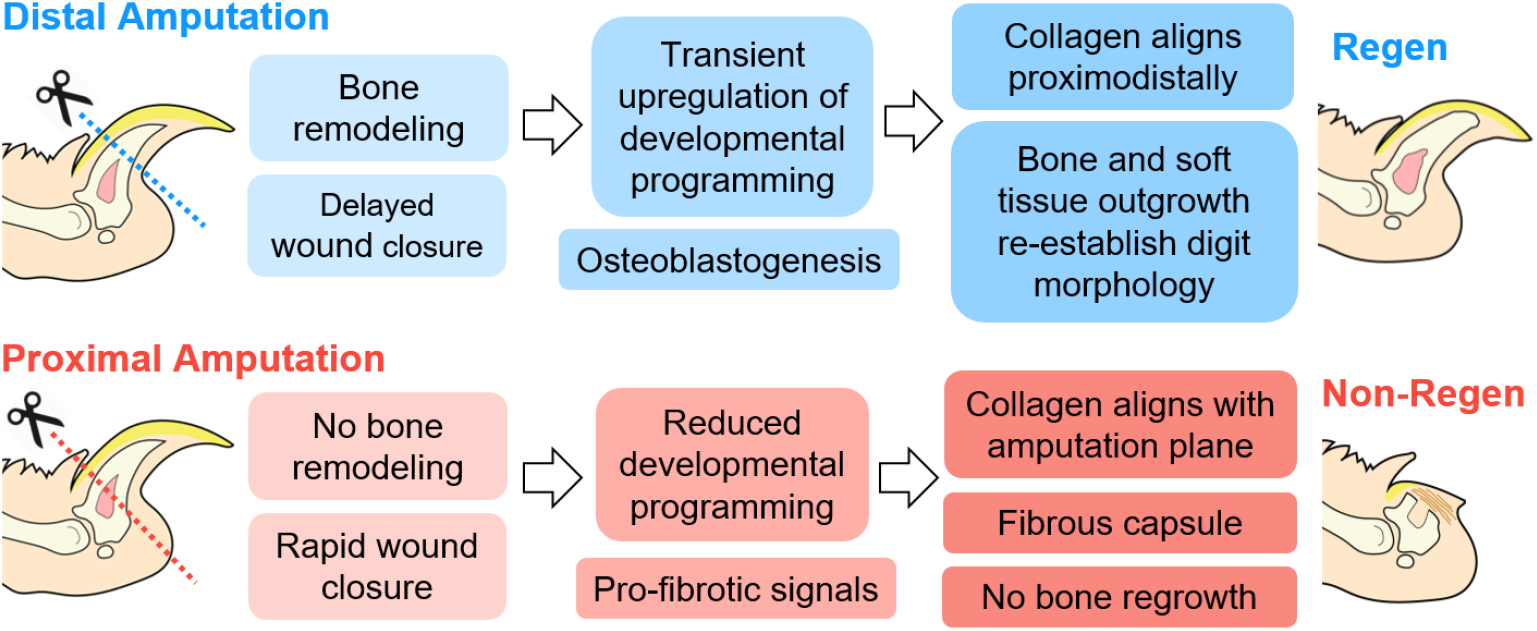
Digit tip regeneration is a level-dependent and spatiotemporally controlled process. After distal amputation of the digit tip, bone degradation is followed by transient upregulation of limb-specific developmental genes and osteogenesis, where the regenerating extracellular matrix becomes aligned in the direction of tissue elongation. In contrast, proximal amputation results in minimal bone remodeling or regrowth. Regenerative failure may be due to reduced developmental signals and cell populations, and/or accelerated fibrosis.

The murine digit represents a unique system to study the factors that lead to regeneration versus scarring, as the extent of the healing process in response to amputation is dependent on the level of resection. We found that a slight difference in P3 amputation level (~0.6 mm more proximal for the Non-Regen group) led to significantly different outcomes. Our transcriptomic analysis, which reflects the resolution of inflammation and bone resorption with the ascension of cell migration, osteoblast differentiation, and bone formation, is consistent with previous reports (Fernando et al., 2011; Simkin et al., 2017; Dawson et al., 2018). Early regeneration is facilitated by delayed wound closure and bone degradation (Simkin et al., 2015; Simkin et al., 2017), events that may promote stem/progenitor cell proliferation and migration from the marrow cavity. In addition, localized ECM degradation releases matrix-bound growth factors, which may stimulate osteoprogenitors (Martin et al., 2009) and/or act as chemoattractants to recruit stem/progenitor cells to the wound site (Agrawal et al., 2011). We demonstrate the expansion of *Hoxa13*+ and *Runx2*+ cells from the open marrow cavity and periosteum, suggesting that the blastema is comprised primarily of mesenchymal-lineage stem/progenitor cells derived from the bone marrow stroma, periosteum, and/or endosteum. In support of this finding, Dawson et al. reported that the blastema is formed by a blend of periosteal and marrow/endosteal cells that differentiate in a progressive wave (defined by *Runx2* and *Sp7* expression) to form new bone distal to the P3 stump (Dawson et al., 2018). Furthermore, removal of the dorsal periosteum at the time of amputation significantly stunted bone regrowth (Dawson et al., 2018), indicating that osteoprogenitors that exist prior to amputation are essential for regeneration. Since these cell populations are reduced by proximal amputation, regenerative failure may be caused in part by an insufficient number of osteoprogenitors. Whether resident osteoblast lineage cells are required for digit regeneration may be queried in future studies using a combination of lineage tracing (Liu et al., 2013) and targeted ablation (Jilka et al., 2009). In addition, digit regeneration is affected by nerve regrowth, which can deliver pro-mitotic growth factors such as fibroblast growth factor-2 (FGF-2) (Takeo et al., 2013) and platelet-derived growth factor-AA (PDGF-AA) (Johnston et al., 2016) to the blastema, as well as influence limb patterning (Rinkevich et al., 2014). Therefore, it is possible that proximally amputated digits fail to establish a hyperproliferative blastema partly due to inadequate peripheral reinnervation and subsequent signaling molecules.

Appropriate spatiotemporal regulation of developmental signaling pathways is essential for organogenesis, especially for the outgrowth and patterning of the limb bud. Limb formation is coordinated by the collinear expression of the HoxA and HoxD complexes (homology groups 9 to 13), a subset of evolutionarily conserved homeobox genes that determine body segment identity (Fromental-Ramain et al., 1996; Villavicencio-Lorini et al., 2010; Zakany and Duboule, 1999). In adults, Hox expression is enriched in mesenchymal stem/progenitor cells in the bone marrow, and is maintained in a distinct, regional pattern that reflects limb development (Rux et al., 2016). The transient upregulation of Hox genes after distal amputation (*Hoxa11/Hoxd11, Hoxa13/Hoxd13*) suggest the activation of a limb-specific developmental pathway concurrent with blastema formation and differentiation, supporting our hypothesis. In addition, we found that *Hoxa13*, a marker of the future autopod (hands and feet) during development, is abundantly expressed by stem/progenitor cells extending from the marrow cavity at 14 DPA. Our analysis of transiently upregulated downstream targets of *Hoxa13*/*Hoxd13* (Hox13) include many of the TFs identified in our regulatory network, including *Prrx1, Alx4, Tbx2*, and *Runx2*. Conversely, *Hoxa13* expression after proximal amputation steadily declined after 12 DPA, consequently reducing the expression of these downstream signals. Knocking out Hox13 leads to the loss of the autopod (Fromental-Ramain et al., 1996), demonstrating that these genes help trigger a digit-specific program. However, since RNA-seq data represent the mean transcript levels of the bulk tissue, it is unclear whether these results are due to dynamic gene expression in single cells, fluctuations in cell populations, or both. To identify the cellular heterogeneity masked by bulk sequencing, single-cell RNA-seq may be utilized to determine which cell types, including rare cell populations, are present throughout regeneration, as well as their varying stages of differentiation (Gerber et al., 2018). Another consideration in the interpretation of our findings is that the transcriptomic landscape may not correspond to protein expression due to post-transcriptional events. Therefore, an integrated analysis of transcriptomic and proteomic data, ideally with single-cell resolution, may provide additional insights into the mechanisms regulating regenerative or non-regenerative responses.

Establishing a provisional matrix that acts as an organizing scaffold is also a crucial step of digit regeneration. After distal amputation, collagen fibers emanate from the degraded P3 bone and orient along the proximodistal axis. Since Hox13 and *Runx2* also direct ECM and collagen fibril organization, their activation may contribute to the aligned collagen microstructure seen in regenerative digits. Indeed, *Hoxd13* may regulate cell polarity, and subsequently collagen deposition, during longitudinal bone growth (Kuss et al., 2014). Marrero and colleagues showed that a reticular fiber-rich scaffold appears during the early stages of regeneration (Marrero et al., 2017), which may provide spatial instruction for stem/progenitor cell migration and differentiation. In contrast, the bone stump of non-regenerative digits becomes circumscribed by a fibrous capsule, reminiscent of the fibrotic scarring observed after amputation of the middle phalanx bone (P2) (Dawson et al., 2017). Well established by 14 DPA, this biophysical barrier may prevent cell proliferation, migration, and subsequent tissue regeneration (Qu et al., 2015). Re-injury of P2 coupled with the delivery of bone morphogenetic protein-2 (BMP-2) can induce bone regrowth if the marrow cavity is exposed, suggesting that regeneration is restricted by the wound microenvironment, rather than the regenerative capacity of endogenous cells (Dawson et al., 2017). Interestingly, we observed higher levels of *H19*, a parentally imprinted lncRNA that is expressed during embryogenesis and repressed postnatally (Gabory et al., 2009), at the bone interface of non-regenerative digits compared to regenerative digits at early time points. Although what causes the epigenetic activation of *H19* in this model is unknown, *H19* is linked to growth suppression by silencing insulin-like growth factor-2 (IGF-2)- dependent signaling (Venkatraman et al., 2013), and its absence results in an overgrowth phenotype (Gabory et al., 2009). In addition, *H19* is a functional regulator of fibrosis and is implicated in the development of keloids and fibrotic diseases of the liver, myocardium, and lung (Lu et al., 2018; Zhang et al., 2018; Zhang et al., 2016). We propose that *H19* expression at early time points may inhibit stem/progenitor cell expansion and instead expedite fibroblast proliferation (Zhang et al., 2016) and type I collagen production (Lu et al., 2018). Alternatively, *H19* may accelerate osteoblast differentiation at the bone interface, reducing the pool of dividing osteoprogenitor cells (Huang et al., 2015). We also found *H19* expressed at the tip of regenerative digits at 21 DPA, suggesting that *H19* may help prevent overgrowth once the regenerate structure is established. Collectively, our data suggest that proximally amputated digits attempt a regenerative response around 12 to 14 DPA but eventually fail, possibly due to insufficient stem/progenitor populations, improper temporal cues, and/or accelerated fibrosis. Therefore, one strategy to stimulate regeneration after proximal amputation may be the exogenous administration of degradative enzymes, which can decrease ECM density and stiffness (Qu et al., 2018) and trigger cell proliferation and migration (Qu et al., 2017; Qu et al., 2015). Modulating early scar formation by targeting fibrotic signaling pathways (Li et al., 2017) may also give stem/progenitor cells an opportunity to expand and differentiate. Finally, spatiotemporal delivery of morphogenic signaling molecules by genetically modified cells and/or biomaterials may help direct and sustain the regenerative process (Dawson et al., 2017; Taghiyar et al., 2017; Qu et al., 2017).

In conclusion, digit regeneration is a complex process that balances both pro-regenerative and pro-fibrotic signaling pathways, making this one of the few *in vivo* mammalian models of musculoskeletal regeneration. Identifying the cellular and molecular mechanisms that regulate regeneration will open new avenues to therapies that will help restore the structure and function of the lost tissues. Furthermore, a better understanding of regeneration failure may shed light on fibrotic diseases and aberrant tissue repair, leading to innovative methods that promote scarless wound healing in adults.

## Materials & Methods

### Digit amputation model

To investigate the effect of amputation level on adult murine digit regeneration, we performed bilateral resection of the digit tips of twenty female C57BL/6J mice at 10–12 weeks old (Jackson Laboratories). This study was carried out in strict accordance with the recommendations in the Guide for the Care and Use of Laboratory Animals of the National Institutes of Health, with all procedures approved by the Institutional Animal Care and Use Committee of Washington University in St. Louis (Protocol #20170244). Mice anesthetized with isoflurane were subjected to amputation of digits 2 and/or 4 of the hind limbs under a dissection microscope and received sustained-release buprenorphine (0.5 mg/kg subcutaneously) for analgesia. Regenerative (Regen) and Non-Regenerative (Non-Regen) states were established by resection of 20–30% (distal amputation) and 60–70% (proximal amputation) of the length of the terminal phalanx bone (P3), respectively. Unamputated digit 3 of the same limb was used as a control. To establish the feasibility of multiple amputations per limb, both single (digit 4 only) and double-digit amputations (digits 2 and 4) were tested (n=2 mice/group/time point). Animals were allowed free cage activity post-operatively until euthanasia via CO2 narcosis at either 10 (wound closure), 12 (blastema formation), 14 (blastema differentiation), 21 (bone regrowth), or 56 days post-amputation (DPA) (bone restoration) in accordance with institutional policy for the humane sacrifice of animals.

### MicroCT imaging and analysis

P3 morphology was assessed with a micro-computed tomography (microCT) scanner designed for *in vivo* use (Bruker SkyScan 1176). Mice anesthetized with isoflurane were placed on the scanner bed in sternal recumbency. Scans of the hind limb digits were obtained at 9 µm resolution (200 µA, 50 kV, 1 mm Al filter, 47° rotation/step) at baseline (pre-amputation; 0 DPA), immediately post-amputation (0 DPA), and after euthanasia, with additional imaging at 28 DPA for 56 DPA animals. Scans were processed using NRecon and DataViewer (Bruker) to obtain sagittal sections of each digit. P3 volume and mid-sagittal length were quantified using CTAn (Bruker) and normalized to pre-amputation values. The mid-sagittal length was defined as the average of measurements from the dorsal and ventral proximal edges of P3 to the distal tip, acquired at the volume centroid. Digits from both single and double-digit models were pooled for statistical analysis (n=4 mice, 6 digits/group/time point).

### Histology and SHG imaging

Harvested digits were fixed in 10% neutral buffered formalin (NBF) at room temperature (RT) overnight. Next, samples were decalcified in 10.6% buffered formic acid at 4°C overnight (Cal-Ex II, Thermo Fisher Scientific) and processed for histologic analysis. Mid-sagittal sections were cut with a paraffin microtome (8 µm thickness) and stained with Hematoxylin and Eosin (H&E) to visualize cells and extracellular matrix (ECM), and with Picrosirius Red (PSR) to visualize collagen (n=3 digits/group/time point). Samples were imaged at 10X magnification with bright field and polarized light microscopy using an inverted light microscope (Olympus VS120) to evaluate collagen deposition and organization (Qu et al., 2015). To visualize the fibrillar collagen microstructure (Qu et al., 2018), unstained paraffin sections were coverslipped using an aqueous mounting medium (VectaMount, Vector Laboratories) for second harmonic generation (SHG) imaging (n=3 digits/group/time point). A tunable coherent Chameleon laser produced an excitation wavelength of 820 nm and SHG signal was collected using a photomultiplier tube detecting wavelengths between 371–442 nm. Tiled *z*-stacks at 1 µm intervals were acquired using a Zeiss LSM 880 with an inverted microscope at 20X magnification (Carl Zeiss Microscopy GmbH), and maximum *z*-stack projections were generated using the open-source platform Fiji.

### RNA isolation, preparation, and sequencing

To evaluate the transcriptomic landscape during digit regeneration versus scarring, bulk RNA sequencing (RNA-seq) was performed on Regen and Non-Regen digits at 12, 14, and 21 DPA (n=4 mice/group/time point). Samples were generated using the double-digit model as described previously, where amputation level and bone regrowth were assessed by microCT. After euthanasia, the tissues distal to the distal interphalangeal joint, excluding the ventral fat pad, were collected under a dissection microscope and stored at −80°C until RNA isolation. Frozen tissues were crushed with a pestle and the tissue lysates homogenized (QIAshredder, QIAGEN). Total RNA was isolated (RNeasy Mini Kit, QIAGEN) (n=4 digits/group/time point) following the manufacturer’s protocol, depleted of ribosomal RNA via a hybridization method (Ribo-Zero Gold rRNA Removal Kit, Illumina), and purified with DNase treatment (Ambion DNA-free DNA Removal Kit, Thermo Fisher Scientific). Library preparation was performed with 1 µg of total RNA per sample (average RNA integrity number of 8), where RNA quality was assessed using an automated electrophoresis system and bioanalyzer (Agilent 4200 Tapestation). Reverse transcription to cDNA was accomplished using SuperScript III RT enzyme (Life Technologies) and random hexamers, with a second strand reaction performed to yield ds-cDNA. cDNA was blunt ended, had an A base added to the 3’ ends, and then had Illumina sequencing adapters ligated to the ends. Ligated fragments were amplified for 11 cycles using primers incorporating unique index tags and then sequenced on an Illumina HiSeq 3000 using single reads extending 50 bases, generating approximately 32 million reads per sample.

### Transcriptomic data analysis

Reads were processed using an in-house pipeline and open-source R packages. Briefly, raw reads were first trimmed using cutadapt to remove low quality bases and reads. Trimmed reads were then aligned to the mouse genome mm10 with GENCODE annotation vM15 using STAR (v2.5.4) with default parameters. Transcript quantification was performed using featureCounts from the subread package (v1.4.6-p4). Further quality control assessments were made using RSeQC and RSEM, and batch-correction was performed using edgeR, EDASeq and RUVSeq. Gene type and transcription factor (TF) annotation were performed using mouse GENCODE vM15 and AnimalTFDB, respectively. Principle component analysis and differential expression analysis for Regen and Non-Regen groups were determined using DESeq2 in negative binomial mode using batch-corrected transcripts from featureCounts (>2-fold expression change, >1 count per million (CPM), Benjamini corrected *P*<0.05) (Love et al., 2014). Pairwise comparisons were made between time points to determine differentially expressed genes (DEGs) within each group. To examine expression patterns, the average reads per kilobase million (RPKM) per time point for Regen (1513 DEGs) and Non-Regen groups (1400 DEGs) were z-score scaled and used for k-means clustering. DEGs were grouped into four clusters (*Upregulation, Transient Upregulation, Transient Downregulation*, and *Downregulation*) and their relative expression over time was plotted using ggplot2. Gene ontology (GO) enrichment analysis was performed using DAVID (v6.8) for DEGs from each cluster (Benjamini corrected *P*<0.05). Next, to compare the expression of Regen-specific DEGs between all groups and time points, the average RPKM from Non-Regen samples was z-score scaled and joined to the Regen samples in their k-means clusters and plotted using pheatmap. TFs that significantly changed expression in the Regen group were used to build a gene regulatory network (GRN) using Cytoscape (v3.0.2).

### ChIP-seq data analysis

To identify potential TF binding targets, *Hoxa13*/*Hoxd13* (Hox13) and *Runx2* chromatin immunoprecipitation followed by sequencing (ChIP-seq) data from mouse E11.5 distal limb cells (Sheth et al., 2016) and mouse MC3T3-E1 pre-osteoblasts and their mature osteoblast progeny (Meyer et al., 2014), respectively, were downloaded from Gene Expression Omnibus (GEO; GSE81358 and GSE41955). Raw fastq files were aligned to the mouse genome mm10 assembly by the Burrows-Wheeler Alignment tool and then processed using methylQA to generate bed and bigwig files. The MACS (v2.0.10) peak caller was used to compare the ChIP-seq signal to a corresponding input control to identify narrow regions of enrichment (peaks) of Hox13 and *Runx2* that passed a q-value threshold of 0.01. ChIP-seq signal density and peak location were further visualized using the WashU Epigenome Browser. ChIP-seq peaks were assigned to the nearest transcription start site of genes using the annotatePeaks.pl function from HOMER2 to determine the binding targets of Hox13 and *Runx2*. GO enrichment analysis was performed as previously described using transiently upregulated Regen genes that were targets of either Hox13, *Runx2*, or both (Benjamini corrected *P*<0.05).

### RNA FISH and image analysis

To validate RNA-seq findings and to visualize spatiotemporal gene expression, single-molecule RNA fluorescence *in situ* hybridization (FISH) was performed for three genes whose levels varied by group and/or time point: *Hoxa13, Runx2*, and *H19*. Regen and Non-Regen digits from 12, 14, and 21 DPA (n=4 digits/group/time point) were harvested, fixed overnight in 10% NBF at RT, and decalcified in 10% ethylenediaminetetraacetic acid (EDTA) for 10 days at 4°C. Samples were processed for paraffin embedding and cut into sagittal sections (5 µm thickness). Sample pre-treatment and RNA probe hybridization, amplification, and signal development were performed using the RNAscope Multiplex Fluorescent Reagent Kit v2 (Advanced Cell Diagnostics) following the manufacturer’s protocol. Probe signals for *Hoxa13,Runx2*, and *H19* were developed using TSA Plus fluorescein, cyanine 5, and cyanine 3 (PerkinElmer), respectively, and the nuclei were counterstained with 4′,6-diamidino-2-phenylindole (DAPI). Samples were imaged with multi-channel confocal microscopy (Zeiss LSM 880). *Z*-stacks were taken at 20X (tiled image) and 63X magnification to capture the entire digit tip and specific regions of interest (e.g., blastema), respectively. To quantify probe signal distribution throughout the digit, the maximum *z*-stack projection for each channel at 20X was converted into a binary image using Fiji. Particles larger than 20 pixels^2^ were excluded from analysis (autofluorescent red blood cells and nail keratin). A density heat map of each probe signal was generated using the open-source platform Bio7 and the R package spatstat, where the intensity value represents the expected number of random points per unit area. Heat maps were overlaid with schematics created using Adobe Illustrator CS2 that depict the boundaries for bone, skin/nail, and underlying connective tissue.

### Statistical analysis

Statistical analyses were performed using GraphPad Prism (v8.0; GraphPad Software, Inc.). Significance was assessed by two-way analysis of variance (ANOVA) with Tukey’s HSD post-hoc tests to compare between groups (*P*<0.05). Primary metrics include bone length, bone volume, and gene expression level (RPKM) as dependent variables, with independent variables of group and time point. Results are presented as mean±s.d. unless specified otherwise.

## Acknowledgments

The authors gratefully acknowledge Mr. Nicholas Benshoff for his technical assistance with surgery, Dr. Christopher Rowland and Mrs. Sara Oswald for their imaging expertise, and the Genome Technology Access Center at Washington University in St. Louis for RNA library preparation and sequencing.

## Data Availability

High-throughput RNA sequencing results are accessible on the GEO database (GSE131078). All relevant data are available from the authors upon request.

